# Sorting of mitochondrial and plastid heteroplasmy in Arabidopsis is extremely rapid and depends on MSH1 activity

**DOI:** 10.1101/2022.03.28.486093

**Authors:** Amanda K Broz, Alexandra Keene, Matheus Fernandes Gyorfy, Mychaela Hodous, Iain G Johnston, Daniel B Sloan

## Abstract

The fate of new mitochondrial and plastid mutations depends on their ability to persist and spread among the numerous organellar genome copies within a cell (heteroplasmy). The extent to which heteroplasmies are transmitted across generations or eliminated through genetic bottlenecks is not well understood in plants, in part because their low mutation rates make these variants so infrequent. Disruption of *MutS Homolog 1* (*MSH1*), a gene involved in plant organellar DNA repair, results in numerous de novo point mutations, which we used to quantitatively track the inheritance of single nucleotide variants in mitochondrial and plastid genomes in Arabidopsis. We found that heteroplasmic sorting (the fixation or loss of a variant) was rapid for both organelles, greatly exceeding rates observed in animals. In *msh1* mutants, plastid variants sorted faster than those in mitochondria and were typically fixed or lost within a single generation. Effective transmission bottleneck sizes for plastids and mitochondria were *N* ≈ 1 and 4, respectively. Restoring MSH1 function further increased the rate of heteroplasmic sorting in mitochondria (*N* ≈ 1.3), potentially due to its hypothesized role in promoting gene conversion as a mechanism of DNA repair, which is expected to homogenize genome copies within a cell. Heteroplasmic sorting also favored GC base pairs. Therefore, recombinational repair and gene conversion in plant organellar genomes can potentially accelerate the elimination of heteroplasmies and bias the outcome of this sorting process.

**Significance statement:** Mitochondria and plastids play essential roles in eukaryotic life; thus, mutations in these organellar genomes can have severe consequences. In animals, early germline sequestration creates genetic “bottlenecks” providing cell-to-cell variance in mitochondrial mutations upon which selection can act. However, the dynamics of organellar mutations in plants and other organisms that lack early germline segregation remain unclear. Here, we show that sorting of mutations in plant organellar genomes proceeds very rapidly – much faster than in animals. In mitochondria, this process is accelerated by MSH1, a gene involved in recombination and repair of organellar genomes. This suggests that in plants, recombinational repair creates cell-to-cell variance in the frequency of organellar mutations, facilitating selection in the absence of a classical germline bottleneck.

## Introduction

In plants, the genetic system is housed in three different compartments: the nucleus, the mitochondria and the plastids. The products of mitochondrial and plastid genomes perform functions critical for cellular metabolism, including oxidative phosphorylation and photosynthesis. Because organellar genomes are present at high cellular copy numbers, multiple alleles can coexist within a cell, a situation known as heteroplasmy. These variants can create opportunities for selfish competition within cells (1–4) and are of great interest because they are often associated with human disease phenotypes, including inherited disorders due to germline transmission and age-related disorders due to heteroplasmies in somatic tissues (5–7). Because of their important health consequences, the dynamics by which de novo mitochondrial point mutations in mammals spread from initially low frequencies to eventually reach fixation (homoplasmy) within a cell have been investigated in detail. Mammalian mitochondrial genomes undergo physical and genetic bottlenecks that increase variance in heteroplasmic alleles among cells, providing a basis for selection (8–10). Bottlenecks in this sense result from a reduction in the effective population size of organellar genomes, which can be due to processes such as drift, preferential organellar DNA amplification, organellar dynamics and/or gene conversion (11–13). The relative size of these bottlenecks can be calculated by comparing variance in heteroplasmic frequencies between mother and progeny (14, 15), and effective mitochondrial bottleneck size (i.e., modelling the heteroplasmy variance generated between generations as a single sampling event) ranges from ∼10-30 segregating units in humans (16–19).

In contrast to the fairly detailed understanding of mitochondrial heteroplasmy in animal systems, there are fundamental gaps in our knowledge of how mitochondrial and plastid mutational variants sort out in plants. Studies of heteroplasmic frequency in plants have mostly been done with species that show biparental organelle inheritance (20–23), presumably because the exceedingly low point mutation rates in plant organellar genomes (24–26) limit the supply of *de novo* mutations to study. In addition, spontaneous organellar mutations that lead to visible phenotypes in plants (i.e., plastid mutations causing chlorosis or variegation) tend to be severe, and heteroplasmic lines can often only be maintained by vegetative propagation (27, 28). These factors have hampered the study of heteroplasmy dynamics in plants, including the extent to which heteroplasmies are transmitted across generations.

The process by which heteroplasmies arise and spread is expected to differ markedly from the accumulation of mutations in the nuclear genome because of three distinguishing characteristics: mode of inheritance, copy number per cell and mutation rate. Organellar genomes show non-Mendelian inheritance, typically characterized by maternal transmission as opposed to the biparental inheritance of nuclear genomes in sexual organisms. Strict uniparental inheritance does not allow for the generation of novel combinations of alleles through recombination and has historically been expected to result in a buildup of deleterious mutations, through a process known as Muller’s ratchet (29, 30). However, the impact of Muller’s ratchet will depend upon the number of genes, which tends to be low in organelle genomes (for example, there are 57 genes in Arabidopsis mitochondria (31)), and the rate of deleterious mutation. In addition, there is accumulating evidence that biparental inheritance of organellar genomes (sometimes referred to as paternal leakage) is more common than previously appreciated (22, 23, 32, 33). Even infrequent biparental inheritance of organellar DNA could represent a pathway for recombination, slowing or preventing the ‘mutational meltdown’ associated with Muller’s ratchet (34). Alternatively, recent theory posits that the bottlenecking associated with uniparental inheritance of organellar genomes may actually provide a benefit by increasing cell-to-cell variability and improving the efficiency of selection at higher organizational levels (35–38). This benefit may explain why uniparental inheritance of organellar genomes has been retained across eukaryotic lineages.

The number of genome copies per cell is another major difference between the nucleus and organelles that impacts the spread of new mutations. In contrast to the single nuclear genome copy that is passed to the next generation in each gamete, cells can contain numerous mitochondria and plastids, and genome copies within each organelle can reach high numbers. As such, a mutation arising in organellar DNA initially constitutes only a single member of a larger population of non-mutant genome copies in the cell. The size of this population is expected to influence the subsequent genetic dynamics. In Arabidopsis, only a few plastids are present in meristematic cells, while over 50 plastids can be found in cells of mature leaves, and the number of plastid genome copies per cell ranges from around 80 in meristematic tissue to >3,000 in mature leaves (39). Mitochondrial genomes are present at ∼50-100 copies in both egg cells and leaf cells of Arabidopsis (40–42). However, in both cell types, the number of mitochondria per cell exceeds 300 suggesting that many or most mitochondria do not contain a full mitochondrial genome copy (41). The number of organellar genome copies in female gametes of plants varies between species but is typically low, not exceeding 100 mitochondrial genome copies (42). In comparison, a single mouse oocyte contains over 200,000 mitochondrial genomes, the result of proliferation from ∼200 mitochondrial DNA copies found in progenitor germ cells of the embryo (8).

Mutation rates also differ among cellular compartments. In land plants, mitochondria have the lowest mutation rate (as inferred from synonymous substitutions), followed by plastids and then nuclei at a ratio of approximately 1:3:10 (24–26). By comparison, mammalian mitochondrial sequence mutation rates greatly exceed those in the nucleus (26). Although plant mitochondria and plastids have low rates of point mutation, their genomes undergo frequent recombination between repeated sequences, resulting in populations of alternative structures (43–45). Thus, reversible structural heteroplasmies can exist in plant mitochondria and have been associated with cytoplasmic male sterility and other types of phenotypic variation (43, 46, 47).

Our previous work found that the nuclear-encoded protein MutS Homolog 1 (MSH1) reduces plant organellar mutation rates (48), in addition to its previously characterized role in suppression of ectopic recombination (49, 50). MSH1 is part of the larger MutS family of genes involved in mismatch repair and has a unique architecture that includes both mismatch recognition and endonuclease domains (51). It has been hypothesized that MSH1 initiates double strand breaks (DSBs) at mismatched or damaged bases to facilitate repair through homologous recombination (48, 52, 53). The high ploidies frequently associated with organelles provide multiple genome copies for homologous recombination which plays a major role in DNA replication and repair processes in plant organellar genomes (44, 54). Gene conversion is a common outcome of DSBs and homologous recombination, resulting in homogenization of genome copies within a cell (44, 54). It has been hypothesized that gene conversion could act as an alternative mechanism to increase cell-to-cell variance in heteroplasmic frequencies in eukaryotic lineages such as plants that may lack a physical bottleneck associated with germline development (37). Thus, we predict the action of *MSH1* and other genes involved in homologous recombination may accelerate the sorting of heteroplasmies.

We previously identified numerous mitochondrial and plastid heteroplasmies in Arabidopsis *msh1* mutant lines and showed that some of these could be transmitted through meristematic and reproductive tissues to subsequent generations (48). Having this unique genetic material provided us the opportunity to study the dynamics of *de novo* heteroplasmies in plants, both within individuals and across generations in *msh1* mutants and wild type backgrounds. We find that heteroplasmic sorting is rapid in plants – particularly in plastids – and that MSH1 function in mitochondria increases the speed of heteroplasmic sorting. For heteroplasmic variants, we also found that GC base pairs preferentially increased in frequency over AT base pairs. These results imply that gene conversion contributes to a high rate of heteroplasmic sorting in plants and potentially biases the outcome of this sorting process.

## Results

### Identification of heteroplasmic variants in msh1 mutants

The *msh1* mutant background afforded us the opportunity to explore the dynamics of heteroplasmy in Arabidopsis. We selected ten high frequency single nucleotide variants (SNVs) resulting from de novo mutations that were identified in organellar DNA pools of *msh1* mutant plants (Table 1). Each SNV was associated with a specific *msh1* family line. All of the SNVs were transitions (GC→AT or AT→GC mutations), and the majority were in intergenic regions. The frequencies for each SNV measured by allele-specific droplet digital PCR (ddPCR) were repeatable and similar to those obtained from sequencing read counts (Table 1).

**Table 1.** Characteristics of SNV targets.

| SNV Target<br>(nucleotide<br>position) | Reference<br>Nucleotide | Variant<br>Nucleotide | Genetic<br>Context | Variant<br>Frequency<br>(sequencing)* | Variant<br>Frequency<br>(ddPCR)* | Heteroplasmy<br>Identified |
| --- | --- | --- | --- | --- | --- | --- |
| Plastid (26553) | A | G | intergenic | 0.076 | 0.0926 (0.0080) | Yes |
| Plastid (29562) | T | C | intergenic | 0.032 | 0.026 | No |
| Plastid (36873) | A | G | intergenic | 0.059 | 0.032 (0.0024) | Yes |
| Plastid (48483) | A | G | intergenic | 0.028 | 0.033 | No |
| Plastid (61599) | A | G | intergenic | 0.068 | 0.090 | No |
| Plastid (72934) | A | G | *psbB* - CDS | 0.023 | 0.017 | No |
| Plastid (118559) | T | C | *ndhG* - CDS | 0.014 | 0.015 (0.0082) | No |
| Mito (91017) | A | G | intergenic | 0.147 | 0.180 | Yes |
| Mito (143184) | T | C | *rrn26* - rRNA | 0.014 | 0.012 | No |
| Mito (334038) | C | T | intergenic | 0.033 | 0.019 (0.0018) | Yes |

\*Variant frequencies are calculated as the count of variant alleles out of the total copy number in the sample. Variants were initially identified from Duplex Sequencing of purified organellar DNA from a pool of ~60 F3 plants (48). These same samples were used to estimate variant frequency with ddPCR. Standard deviation for ddPCR experiments is shown in parentheses for the SNVs used in subsequent studies, and seven to ten technical replicates were used for calculation of standard deviations.

We used this ddPCR assay to identify heteroplasmy in individual plants by screening young leaves from the siblings and/or progeny of the plants used in the initial identification of SNVs (48). After adjusting for organellar genome copy number and numts (see Methods), only four of the ten SNVs were found to be heteroplasmic in any of the screened individuals; the other six variants had either reached fixation or were not detectable in the individuals screened (Table 1, Table S1). Therefore, we proceeded with these four heteroplasmic SNVs (two mitochondrial and two plastid) for all subsequent experiments and analyses. The number of heteroplasmic individuals identified was higher for the mitochondrial SNVs tested (46.5% of individuals) versus those from the plastid (3.7%) (Table S1), which is why a greater number of markers and total plants were screened for plastid versus mitochondrial SNVs.

### Intergenerational heteroplasmic sorting occurs more rapidly in plastids than mitochondria in an msh1 mutant background

To determine the frequency with which heteroplasmies are transmitted across generations, we used *msh1* individuals that were each heteroplasmic for one of the four identified SNVs as mothers to generate selfed progeny. We expected that variant frequencies in the mother would influence the distribution of heteroplasmy in the progeny, so we used mothers with a wide range of starting allele frequencies. Both mitochondria and plastid SNVs showed rapid sorting over a single generation, as many progeny were fixed for either the wild type or alternative (SNV) allele (Figure 1, Table S2). This trend was particularly striking in plastids: very few heteroplasmic progeny were identified for plastid SNVs, regardless of the heteroplasmic frequency in the mother. For mitochondrial SNVs, progeny showed a tighter and more continuous distribution of heteroplasmy roughly distributed around the allele frequency of the mother, with many progeny retaining a heteroplasmic state.

**Figure 1.**
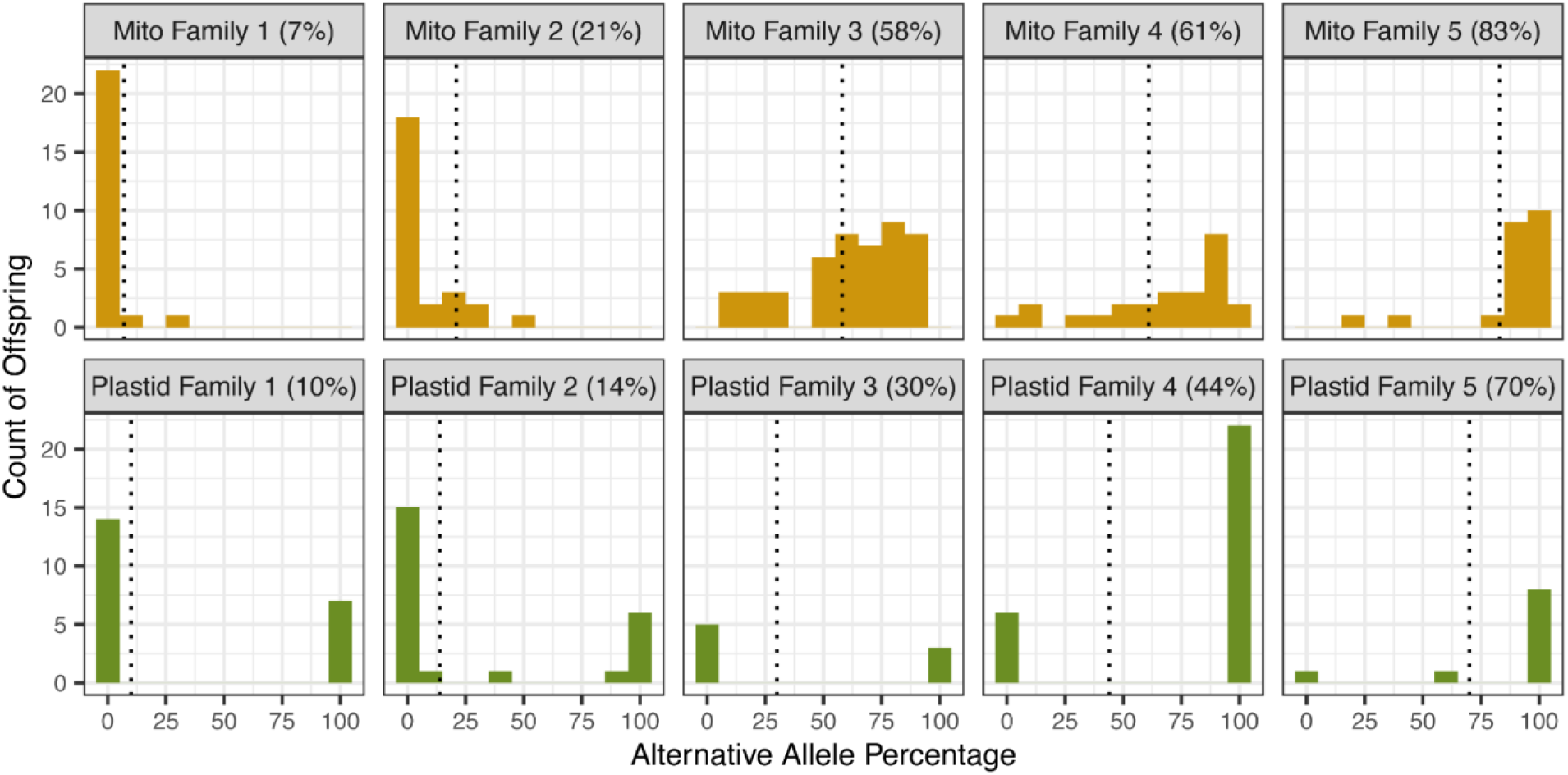
Distribution of heteroplasmy across generations in an *msh1* mutant background. Heteroplasmy (alternative allele frequency) was evaluated using ddPCR of leaf tissue in progeny of maternal lines with different levels of heteroplasmy, indicated by dotted vertical lines and numbers in parentheses on each graph. Histograms indicate the number of individuals showing different levels of heteroplasmy. Mitochondrial families (orange) are depicted in the top row, plastid families (green) are shown on the bottom row. For the sake of comparison with the five mitochondrial families, data for plastid families 6 and 7 in not shown but is included in Table 2.

**Table 2.**
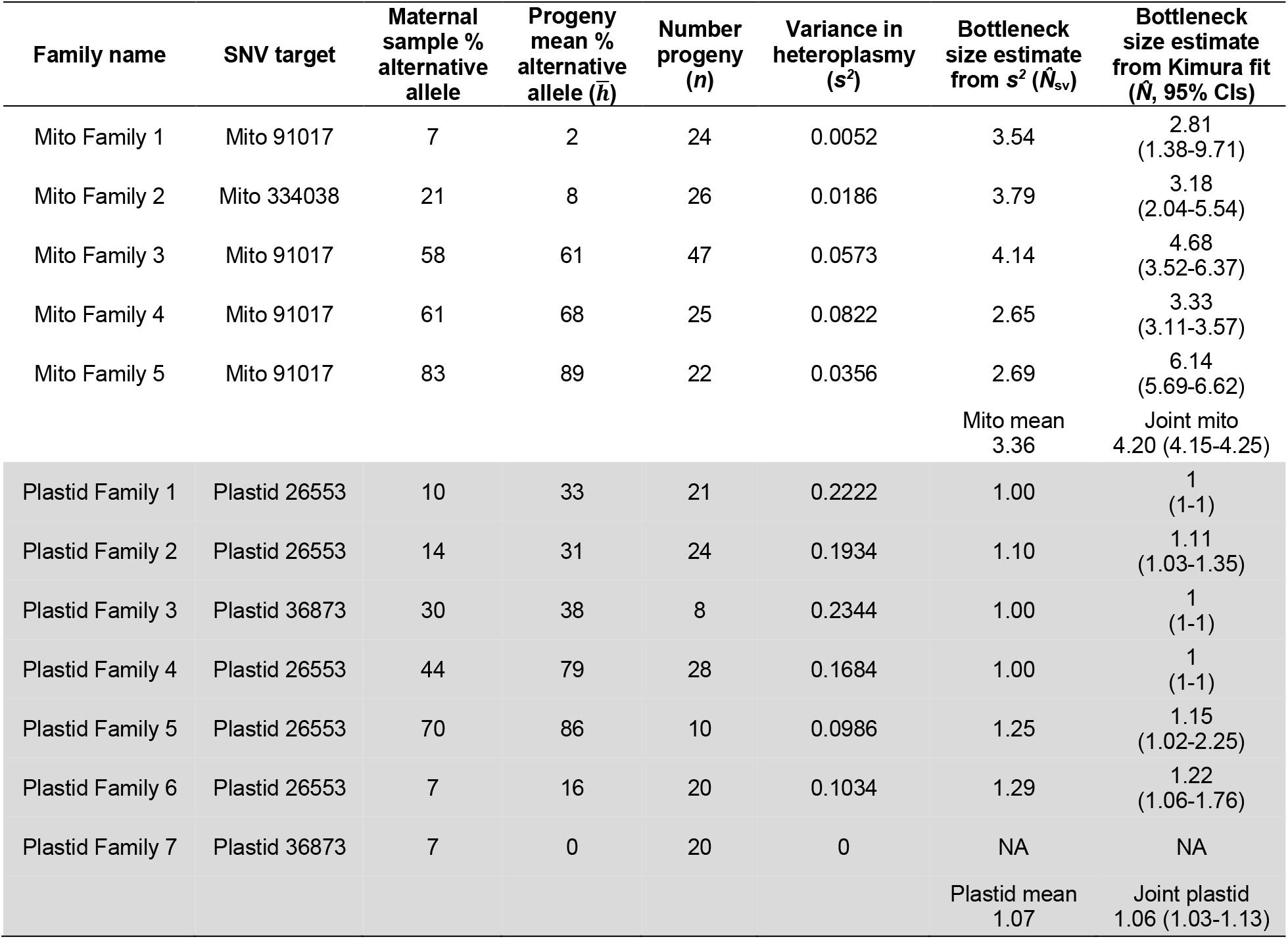
Effective transmission bottlenecks in organellar genomes in the *msh1* mutant background calculated for plastid and mitochondrial markers over single generations in *msh1* mutant plants. Confidence intervals for Kimura estimates are noted in parentheses below calculated bottleneck value. Likelihood ratio test (see methods) show that plastid bottlenecks are significantly smaller than mitochondrial bottlenecks (*p* = 3.2 × 10^-55^).

The distribution of heteroplasmy in progeny derived from a heteroplasmic mother can be used to calculate an effective transmission bottleneck size (*N*). Although this measure has often been (incorrectly) equated with the number of organellar genomes transmitted to the next generation, it is more appropriately thought of as a relative metric to compare the cumulative biological sampling variance in genome transmission throughout development and across generations (14, 15). A common way of estimating bottleneck size is by taking the reciprocal normalized sample variance 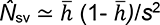, as an estimate for the number of effective segregating units (14), where *h̄* is the sample mean heteroplasmy and *s*^2^ is the sample variance in heteroplasmic frequencies for a set of tissue or progeny samples. However, the use of this statistic is problematic for moderate samples of highly segregated cases, where the sample variance approach yields estimates and uncertainties that are incompatible with the concept of segregating units (for example, inferring *N̂*_sv_ < 1 with confidence intervals that include zero) (55). We took two approaches here to estimate bottleneck size. First, for comparison with previous work, we used the above equation to calculate *N̂*_sv_, but we employed the population variance (without Bessel’s correction) for *s*^2^. This approach yields a slightly biased estimate but one that respects heteroplasmy constraints (*N̂*_sv_ ≥ 1). More rigorously, we also used a maximum likelihood approach based on the Kimura distribution (55) that captures these constraints in order to identify the most likely value (*N̂*) of the bottleneck parameter as well as the confidence interval (CI) of this measure given a full set of observations. We also used this Kimura-based approach to perform hypothesis testing (see Methods). Both values (*N̂*_sv_ and *N̂*) are reported in tables for comparison, but we refer to the more rigorous Kimura estimates in the text.

Using data from seven plastid families and five mitochondrial families we found that, over a single generation, plastids typically have a smaller transmission bottleneck size than mitochondria (Table 2). Plastid transmission bottlenecks were extreme, approximating a value of one (*N̂* = 1.06, 95% CIs: 1.03-1.13), in accordance with observations that the vast majority of offspring were homoplasmic for one allele or the other (Figure 1). The average mitochondrial transmission bottleneck size in *msh1* mutants (*N̂* = 4.20, 95% CIs: 4.15-4.25) was significantly larger than in plastids (*p* = 3.2 × 10^-55^, likelihood ratio test, Table 2). Therefore, the number of effective genome copies passed throughout development and from mother to progeny is larger for mitochondria than for plastids. However, the transmission bottleneck values for both organelles reflect relatively rapid heteroplasmic sorting.

### Heteroplasmic sorting in vegetative and floral tissues occurs more rapidly in plastids than mitochondria in an msh1 mutant background

Although reproductive tissue is one location where heteroplasmic sorting can happen, it can also occur within cells and tissues as they grow. This may be particularly important in organisms like plants that do not exhibit early germline specification (56, 57), a hypothesis that is supported by recent theory predicting ongoing segregation in plant tissues (37). We sampled multiple leaf and inflorescence tissues from selected progeny of *msh1* heteroplasmic mothers and quantified heteroplasmic rates (Figure 2, Table S3). Two plastid family lines (six individuals) initially selected for analysis showed no within-plant heteroplasmy and were either fixed for the wild-type or alternative SNV allele. The other family line showed high variance in heteroplasmy rates across tissues, indicative of rapid within-plant sorting. Using rates of heteroplasmy in the tissue samples from each plant, we found that within-plant bottleneck sizes were significantly smaller on average for plastids (*N̂* = 5.92, 95% CIs: 3.85-9.51) than for mitochondria (*N̂* = 12.00, CIs: 11.95-12.05, *p* = 3.4 × 10^-29^, likelihood ratio test; Table 3). Within-plant bottleneck size was larger than the transmission bottleneck size seen between generations (plastids, *p* = 2.4227 x 10^-29^; mitochondria, *p* = 1.6 × 10^-30^, likelihood ratio tests), suggesting that sorting during vegetative growth indeed contributes to heteroplasmic variance, but the full reproductive cycle and transmission to the next generation involves a tighter effective bottleneck.

**Figure 2.**
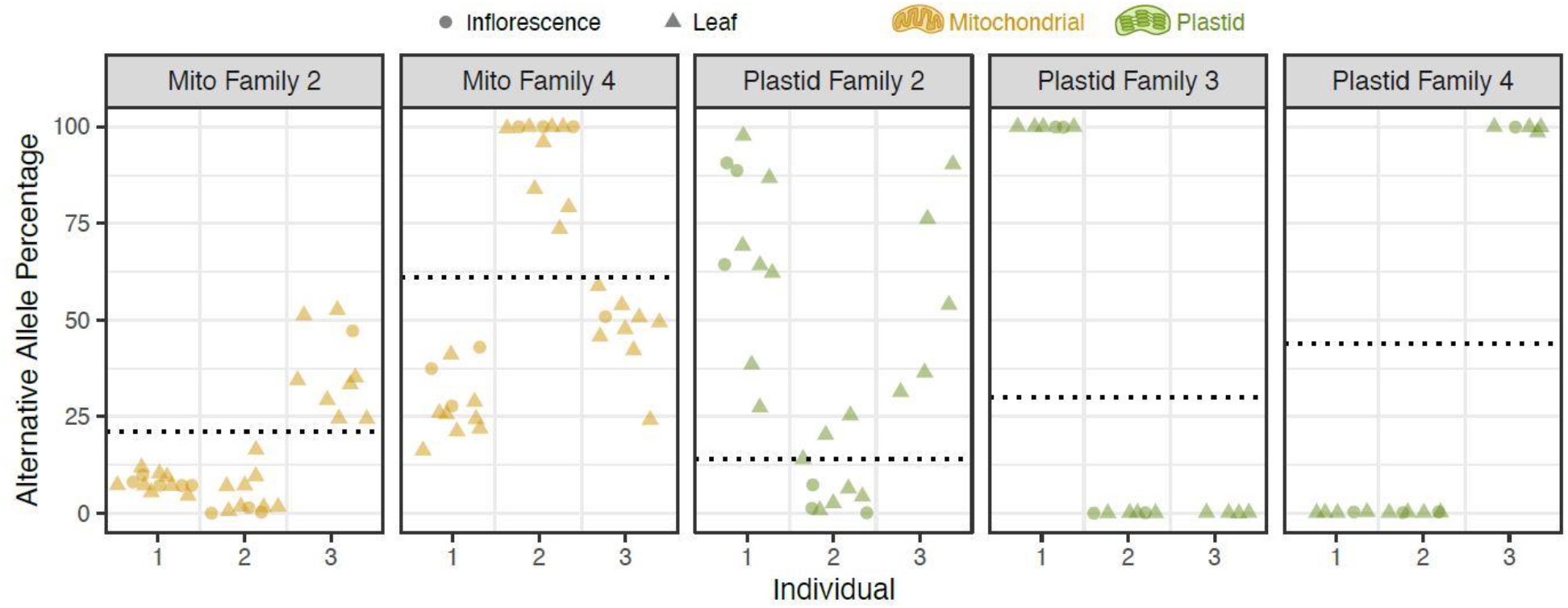
Distribution of heteroplasmy within individual plants in an *msh1* background. Progeny from mothers with varying levels of heteroplasmy (experiment shown in Figure 1) were selected for further analysis. Dotted lines indicate levels of maternal heteroplasmy. Levels of heteroplasmy (alternative allele percentage) for leaf and inflorescence tissues were determined using ddPCR.

**Table 3.**
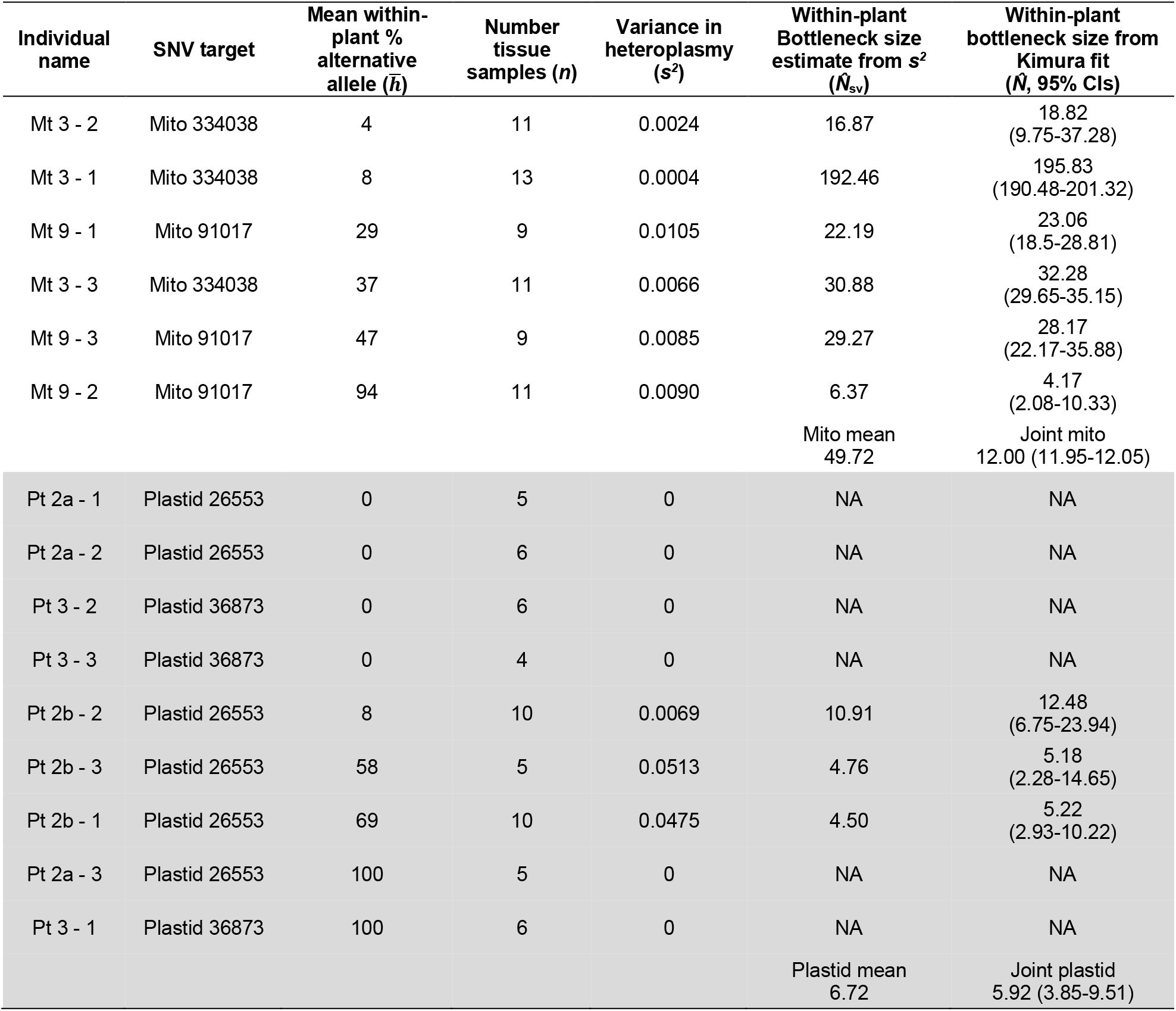
Within-plant bottlenecks in organellar genome transmission in *msh1* mutant backgrounds. Confidence intervals for Kimura estimates are noted in parentheses below calculated bottleneck value. Not applicable (NA) indicates that there was no variation within the tissues tested. Likelihood ratio test (see methods) show that within plant plastid bottlenecks are significantly smaller than mitochondria bottlenecks (*p* = 3.4 × 10^-29^).

| Individual name | SNV target | Mean within-plant % alternative allele ($\bar{h}$) | Number tissue samples ($n$) | Variance in heteroplasmy ($s^2$) | Within-plant Bottleneck size estimate from $s^2$ ($\hat{N}_{sv}$) | Within-plant bottleneck size from Kimura fit ($\hat{N}$, 95% CIs) |
| --- | --- | --- | --- | --- | --- | --- |
| Mt 3 - 2 | Mito 334038 | 4 | 11 | 0.0024 | 16.87 | 18.82<br>(9.75-37.28) |
| Mt 3 - 1 | Mito 334038 | 8 | 13 | 0.0004 | 192.46 | 195.83<br>(190.48-201.32) |
| Mt 9 - 1 | Mito 91017 | 29 | 9 | 0.0105 | 22.19 | 23.06<br>(18.5-28.81) |
| Mt 3 - 3 | Mito 334038 | 37 | 11 | 0.0066 | 30.88 | 32.28<br>(29.65-35.15) |
| Mt 9 - 3 | Mito 91017 | 47 | 9 | 0.0085 | 29.27 | 28.17<br>(22.17-35.88) |
| Mt 9 - 2 | Mito 91017 | 94 | 11 | 0.0090 | 6.37 | 4.17<br>(2.08-10.33) |
|  |  |  |  |  | Mito mean<br>49.72 | Joint mito<br>12.00 (11.95-12.05) |
| Pt 2a - 1 | Plastid 26553 | 0 | 5 | 0 | NA | NA |
| Pt 2a - 2 | Plastid 26553 | 0 | 6 | 0 | NA | NA |
| Pt 3 - 2 | Plastid 36873 | 0 | 6 | 0 | NA | NA |
| Pt 3 - 3 | Plastid 36873 | 0 | 4 | 0 | NA | NA |
| Pt 2b - 2 | Plastid 26553 | 8 | 10 | 0.0069 | 10.91 | 12.48<br>(6.75-23.94) |
| Pt 2b - 3 | Plastid 26553 | 58 | 5 | 0.0513 | 4.76 | 5.18<br>(2.28-14.65) |
| Pt 2b - 1 | Plastid 26553 | 69 | 10 | 0.0475 | 4.50 | 5.22<br>(2.93-10.22) |
| Pt 2a - 3 | Plastid 26553 | 100 | 5 | 0 | NA | NA |
| Pt 3 - 1 | Plastid 36873 | 100 | 6 | 0 | NA | NA |
|  |  |  |  |  | Plastid mean<br>6.72 | Joint plastid<br>5.92 (3.85-9.51) |

### MSH1 activity accelerates heteroplasmic sorting in mitochondria

Because MSH1 is hypothesized to introduce DSBs and promote recombinational repair (48, 52, 53), we predicted that a functional copy of the *MSH1* gene would speed up heteroplasmic sorting by homogenizing genome copies through gene conversion, as predicted by theoretical modeling (37). To test this hypothesis, we transferred heteroplasmic variants to a wild-type background by crossing heteroplasmic *msh1* female plants with wild-type males and analyzing heteroplasmy levels in both F1 and F2 (selfed) progeny. All plants were genotyped at the *MSH1* locus, and only individuals that were heterozygous or homozygous wild type were included in the heteroplasmy analysis (the *msh1* mutation is recessive). Due to the extremely low number of individuals that were heteroplasmic for plastid SNVs and the rapid plastid heteroplasmic sorting rates, this backcrossing method was only successful in generating lines to study mitochondrial heteroplasmy. For mitochondrial SNVs in the wild-type background, we saw extremely rapid sorting of heteroplasmies (Figure 3, Table S2, Table S4), akin to our observations in plastids under *msh1* mutant backgrounds (Figure 1). The average mitochondrial transmission bottleneck size for wild-type plants (*N̂* = 1.33, 95% CIs: 1.20-1.54) was significantly reduced from that in the *msh1* mutant background (*N̂* = 4.20, 95% CIs: 4.15-4.25, *p* = 8.1 × 10^-49^, likelihood ratio test; Table 2, Table 4). The standing number of organellar genomes per nuclear genome copy in leaf tissue did not differ significantly between *msh1* and wild type backgrounds (Figure S1), suggesting that the differences we identified in heteroplasmic sorting were more likely due to differences in MSH1 activity rather than changes in the physical number of organellar DNA copies per cell.

**Figure 3.**
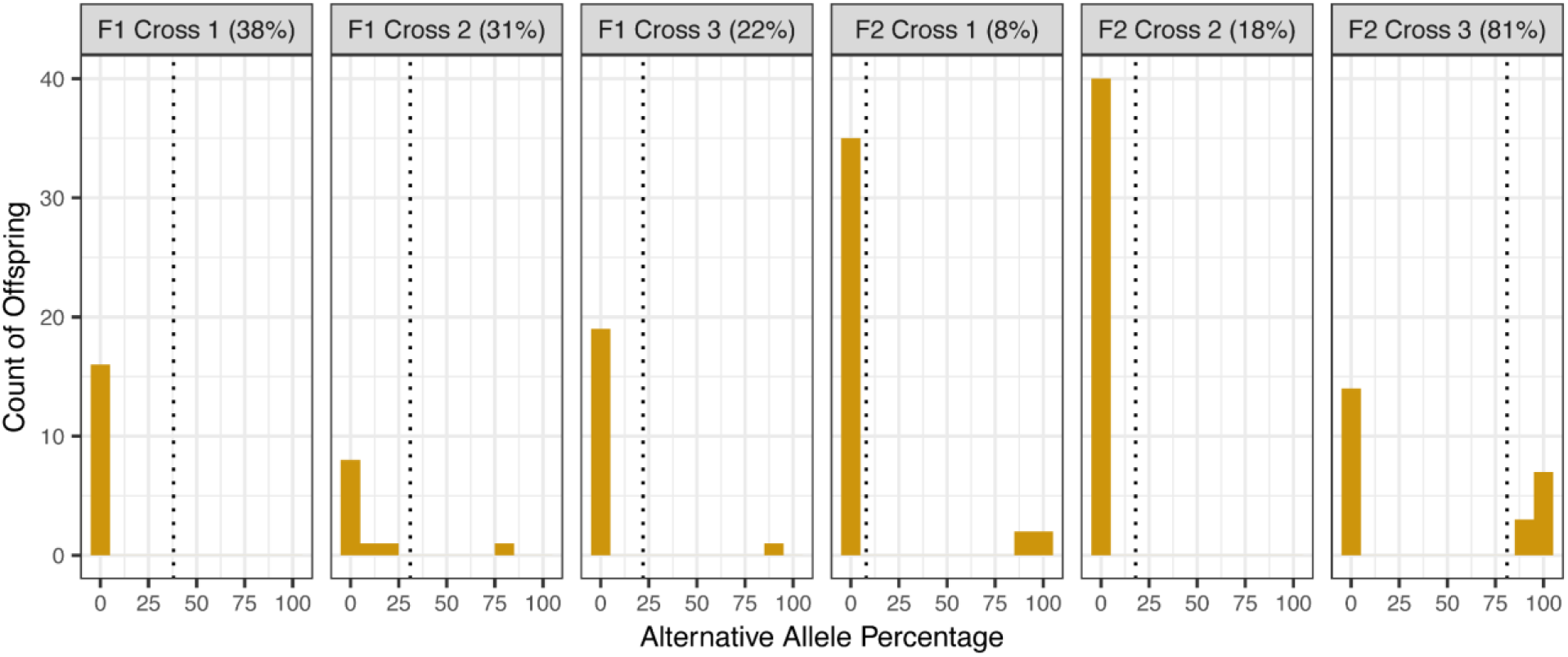
Distribution of mitochondrial heteroplasmy across generations in a wild-type background. Mitochondrial heteroplasmies were backcrossed into a wild-type *MSH1* background. Heteroplasmy (alternative allele frequency) was evaluated using ddPCR of leaf tissue in progeny of maternal lines with different levels of heteroplasmy, indicated by vertical dotted lines and numbers in parentheses on each graph. Histograms indicate the number of individuals showing different levels of heteroplasmy. All three F2 mothers were progeny from F1 Cross 2.

**Table 4.** Effective transmission bottlenecks in organellar genomes in wild type background. Effective transmission bottleneck size was calculated for mitochondrial markers over single generations in wild type plants. Confidence intervals for Kimura estimates are noted in parentheses below calculated bottleneck value. Not applicable (NA) indicates that there was no variation within the progeny – they were all fixed for one allele or the other. Note that bottleneck size estimates for F1 cross 1 are imprecise because they are based on a family in which only one of 17 progeny remained heteroplasmic and that individual had a low allele frequency (2%). The large point estimates for that family weigh heavily when taking a mean of *N̂*_sv_ estimates but do not bias the joint maximum-likelihood estimate of *N̂*.

| Cross type | SNV target | Maternal sample % alternative allele | Progeny mean % alternative allele ($\bar{h}$) | Number progeny ($n$) | Variance in heteroplasmy ($s^2$) | Bottleneck size estimate from $s^2$ ($\hat{N}_{sv}$) | Bottleneck size estimate from Kimura fit ($\hat{N}$, 95% CIs) |
| --- | --- | --- | --- | --- | --- | --- | --- |
| F1 cross 1 | Mito 334038 | 38 | 0.14 | 17 | 0.00003 | 46.13 | 18.18<br>(2.45-204.07) |
| F1 cross 2 | Mito 334038 | 31 | 8 | 10 | 0.05867 | 1.63 | 1.94<br>(1.26-4.45) |
| F1 cross 3 | Mito 91017 | 22 | 5 | 20 | 0.03708 | 1.15 | 1.58<br>(1.13-3.67) |
| F2 cross 1 | Mito 334038 | 8 | 18 | 32 | 0.13963 | 1.05 | 1.26<br>(1.11-1.62) |
| F2 cross 2 | Mito 334038 | 18 | 0 | 37 | 0.00000 | NA | NA |
| F2 cross 3 | Mito 334038 | 81 | 36 | 24 | 0.21511 | 1.07 | 1.24<br>(1.11-1.54) |
|  |  |  |  |  |  | Mean<br>10.21 | Joint estimate<br>1.33 (1.20-1.54) |

### GC-biased inheritance in organellar genomes

The fact that the SNVs used in our analysis were GC→AT or AT→GC transitions created the opportunity to search for GC or AT bias in the inheritance of SNVs in plant organellar genomes. Using the allele frequency data from our heteroplasmic mothers and progeny, we found that, even though angiosperm organelle genomes are typically AT rich (58), there is evidence of a GC bias during heteroplasmic sorting in both plastids and mitochondria (Figure 4, Table S5). The frequency of the GC allele increased in the progeny relative to the mother in 14 of 17 families (two-sided binomial test, *p* = 0.0127). Although modest increases in the frequency of the GC allele were found in most lines, five showed that mean progeny GC frequency increased over 20% compared to the mother (Figure 4B). The mean increase in frequency for the GC allele was 14.1%, which differed significantly from zero (two-sided t-test, *p* = 0.0039). The magnitude of this increase was nearly identical for mitochondrial SNVs (14.0%) and plastid SNVs (14.2%).

**Figure 4.**
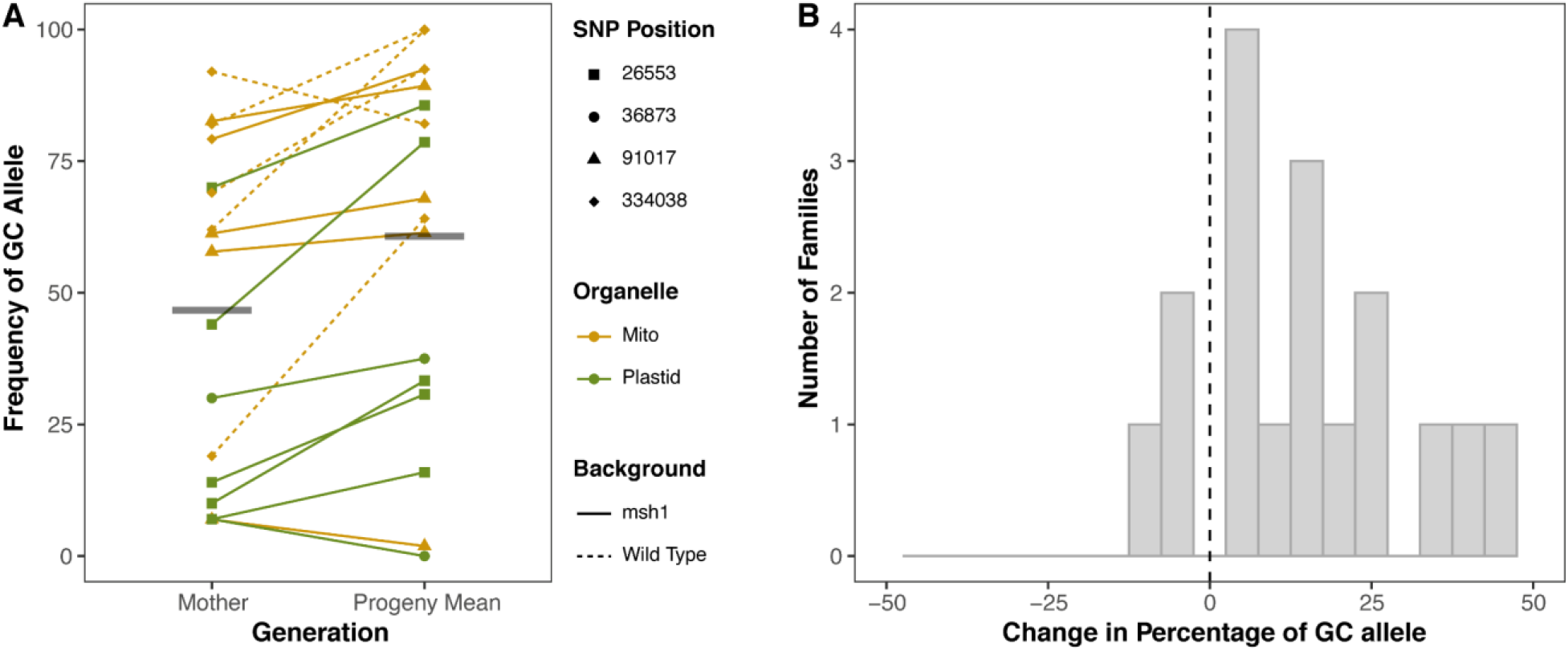
Changes in frequency of GC alleles between generations. A) Norm of reaction plot showing the frequency of GC alleles (the SNVs from this study) in mothers and progeny (mean value). The reference (wild type) allele was AT and the variant GC in all cases except for mt334038 in which the inverse was true (Table 1). Means for each group are shown with horizontal gray bars. B) Histogram showing the change in percentage of GC alleles for tested family lines. A two-sided t-test shows bias toward GC alleles, *p* = 0.0039.

## Discussion

### Potential causes of rapid heteroplasmic sorting in plant organelles

In Arabidopsis, we found that both plastid and mitochondrial heteroplasmies sorted out to homoplasmy within one to a few generations and exhibited tight effective bottlenecks, regardless of whether they were found in wild type or *msh1* mutant backgrounds (*msh1* plastid *N̂* ≈ 1, *msh1* mitochondria *N̂* ≈ 4, wild-type mitochondria *N̂* ≈ 1.3). In contrast, effective mitochondrial germline bottlenecks are estimated to be substantially larger in most animal systems: ∼5-10 in *Daphnia* (59), ∼9 in macaques (60), ∼10-30 in humans (16–19), ∼30 in *Drosophila* (61), ∼60 in *Caenorhabditis elegans* (62), ∼80 in salmon (63), 5-100 in mice (9, 55), and 170 in zebrafish (64). Thus, in animals, heteroplasmy is often retained over multiple generations.

Germline bottlenecks are well established as drivers of heteroplasmic variance in animals, but the mechanism is still not completely understood (8, 9, 14, 18). The bottleneck is thought to be due, at least in part, to the physical reduction of mitochondrial DNA copies during female germline development (65, 66). However, other lines of evidence suggest that selection (67–69), mitochondrial dynamics (12), and preferential genome amplification (11, 13) can also play roles in reducing the effective population size of mitochondrial genomes.

In plants, where the germline is segregated later in development (56, 57, 70), the way in which heteroplasmic variance increases is even less clear. However, the relatively modest number of mitochondrial and plastid genome copies in meristematic and reproductive tissues is a likely contributor to rapid heteroplasmic sorting. In plants, all aboveground tissues, including reproductive organs, are derived from the shoot apical meristem (SAM). Arabidopsis SAM cells contain only ∼80 copies of plastid DNA housed within 4-10 proplastids (39). The number of mitochondrial genome copies in SAM cells is not well established, but these values are typically <100 in both vegetative tissues and egg cells (40–42). Additional mechanisms, including selection could reduce the effective population size of organelle genomes by acting as an effective germline bottleneck. The frequent recombination associated with plant organellar genomes (44, 54) could also act as an effective bottleneck by homogenizing genomes through gene conversion without requiring a physical reduction in organelle DNA copy number, as predicted by recent theory (37). The rapid heteroplasmic sorting in Arabidopsis organelles, as well as our finding that MSH1 activity further accelerates the rate of sorting (see below), supports this hypothesis.

Although the rate of heteroplasmic sorting has not been studied extensively in plants, there are supporting lines of evidence that this process may be rapid in other angiosperms. For example, controlled crosses of *Silene vulgaris* and *Daucus carota* have shown that mitochondrial heteroplasmy is maternally transmitted at only low levels across generations (20, 21). Mitochondrial variants produced by repeat-mediated recombination have been observed to rapidly rise from mean cellular copy numbers under one to high frequencies or even homoplasmy across cells, through a process called substoichiometric shifting (SSS) (43, 46). Because of the role of *MSH1* in recombination surveillance, these structural variants arise more frequently in *msh1* mutants (50), but the causes of rapid SSS have remained less clear. Studies of variegation mutants derived from biparental plastid inheritance show that sorting occurs rapidly at the level of whole organelles and is frequently complete within a single generation (15, 28, 71). When plastid DNA is modified using a transgenic approach, homoplasmy is typically reached after a few rounds of antibiotic selection (72, 73). The rapid sorting of transgenic plastid mutations has also been observed in the absence of antibiotics (74). It is currently unclear whether rapid sorting of plastid transgenes occurs in all plants as it has been extremely difficult to obtain homoplasmic transgenic lines of various species, particularly monocots (75–77). However, this may be due to the general recalcitrance of these species to plastid transformation or inefficient selection that makes variants difficult to detect (75, 76). The approach we have used here would be valuable to test heteroplasmic sorting in other species because most intergenic SNVs are unlikely to be under strong selective pressure, as opposed to the entire genes that are introduced with transgenic approaches or the large structural rearrangements associated with SSS. The rapid sorting we have identified could readily explain these diverse instances of segregation and occasional amplification of rare variants in organellar genomes, but characterization from diverse species is needed because the developmental and genetic mechanisms of heteroplasmic sorting may vary across plant lineages. However, the elevated germline expression of recombination machinery including MSH1 appears to be conserved across several angiosperms (37), suggesting a similar genetic basis may exist for generating heteroplasmic variance.

### Differences in heteroplasmic sorting between mitochondria and plastids

In the *msh1* mutant background, we found that plastids had tighter bottlenecks than mitochondria, both across generations (*N̂* = ∼1 versus ∼4, respectively) and within individuals (*N̂* = ∼6 versus ∼12, respectively). This result may seem surprising because the much greater relative copy number of plastid versus mitochondrial DNA typically seen in most aboveground tissues (39, 41) would be expected to result in lower levels of heteroplasmic variance in plastids (i.e., a wider bottleneck size). However, as noted above, mitochondria and plastids both possess low genome copy numbers in the SAM and/or reproductive tissues (39–42), presenting similar potential for physical bottlenecks. One key difference between the two types of organelles is that mitochondria can experience both full and transient fusion events over the plant life cycle allowing for genetic exchange, whereas plastids rarely if ever fuse (15, 78–82). Mitochondrial fusion is particularly prevalent in the SAM, where it is estimated that 80% of the mitochondrial volume is fused into a dynamic tentaculate cage-like structure, creating the opportunity for sharing of genome copies between formerly distinct mitochondrial compartments (79). Here, gene conversion is predicted to homogenize the mitochondrial genomes within a cell leading to increased heteroplasmic variance between cells (37, 79, 83). Thus, all else being equal, existing theory would predict that fusion results in more rapid heteroplasmic sorting in mitochondria than plastids – the opposite of what we observed. This suggests that other differences in organelle biology can impact the rate of variability in organellar DNA populations.

Additional factors are expected to influence the speed of heteroplasmic sorting, but the extent to which they differ between plastids and mitochondria is often unclear. These processes include the number of organelles per cell, rates of organellar DNA replication and degradation, rates of organelle turnover, the physical partitioning of organellar DNA during organelle replication, and the partitioning of whole organelles during cell division (15). In addition, plastids and mitochondria house distinct versions of many DNA replication, recombination and repair proteins (54), which could influence relative rates of gene conversion. It is also possible that dual-targeted proteins involved in these pathways, such as MSH1, may differentially impact gene conversion rates between organelles. All of these factors likely vary based on cell type and developmental stage, and they may work in combination. Models of heteroplasmic sorting have tried to integrate many of these factors, but much of the relevant biological data required for parameterization is still lacking, particularly in plant systems (9, 14, 18, 37, 84).

### The role of MSH1 in heteroplasmic sorting

We found that mitochondrial heteroplasmic sorting was faster in wild type Arabidopsis plants than in *msh1* mutants (*N̂* = ∼1.3 versus ∼4 respectively), with the majority of wild type progeny reaching fixation for either the reference or alternative allele within a single generation. This result lends support to the hypothesized repair mechanism in which MSH1 identifies mismatches and initiates DSBs followed by template-based recombinational repair [(48, 52, 53), Figure 5A]. Under this model, MSH1 would increase rates of gene conversion, thereby homogenizing genome copies within cells and increasing variance among cells [(37), Figure 5B]. Somewhat confusingly, MSH1 is primarily known as a recombination suppressor because it performs organellar genome surveillance, preventing *illegitimate* recombination between small repeats that can result in genome rearrangements and instability (49, 50). However, this role is not contradictory to the hypothesis that MSH1 activity increases the overall rate of *homologous* recombination. Indeed, these patterns may well reflect the same mechanism of action, in which MSH1 promotes homologous recombination by introducing DSBs at any mismatched bases, regardless of whether they were generated by strand invasion between short/imperfect repeats or by DNA replication errors. Although we might predict that this same mechanism would increase heteroplasmic sorting rates in plastids, we were unable to generate wild type plants that were heteroplasmic for plastid markers – thus the role of MSH1 in plastid heteroplasmic sorting remains an open question.

**Figure 5.**
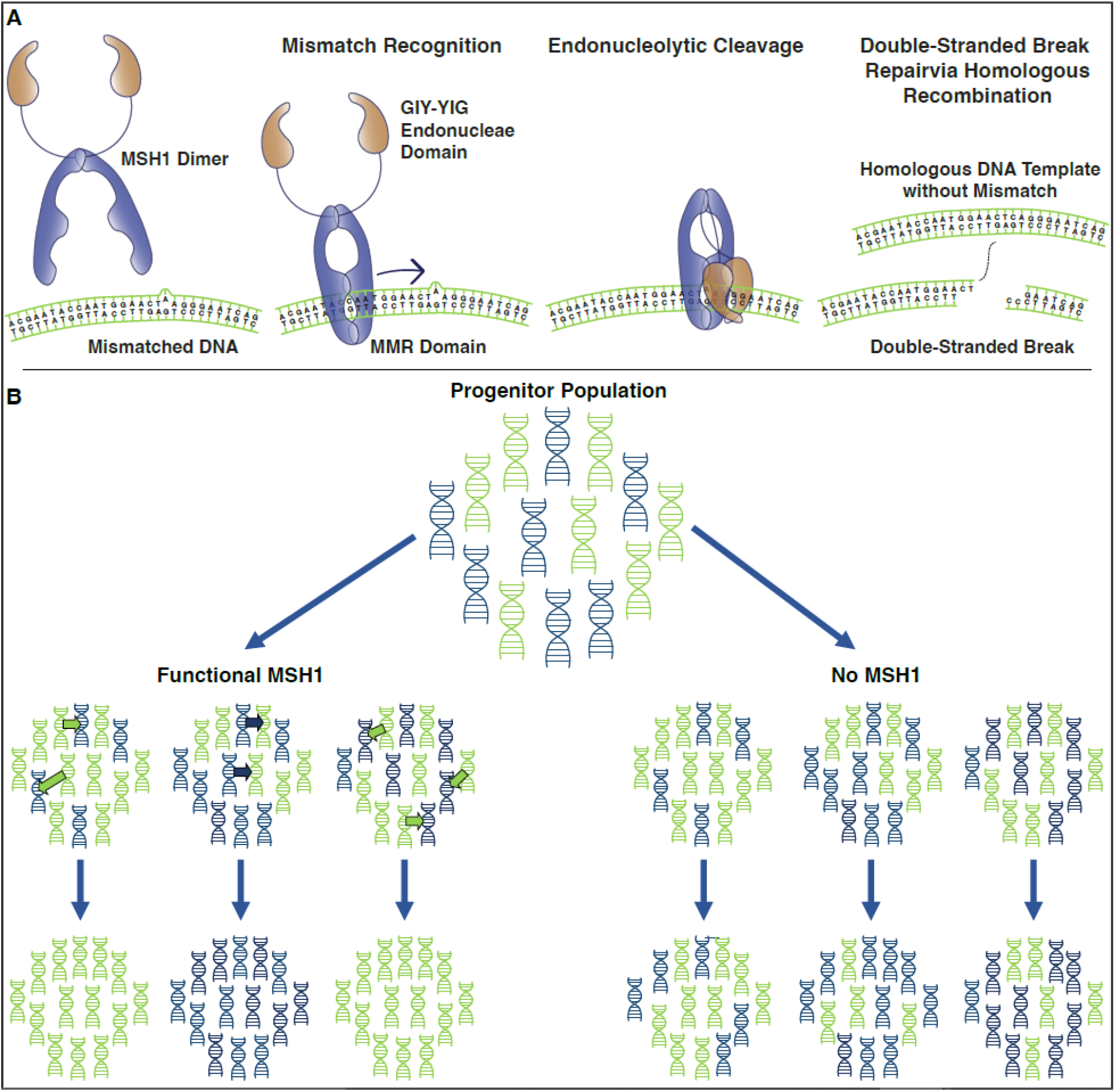
Hypothesized model for role of MSH1 in accelerating heteroplasmic sorting. A) Previously hypothesized mechanism (48, 52, 53) through which MSH1 promotes homologous recombination and repair. The mismatch repair (MMR) domain of dimeric MSH1 slides along organellar DNA until it reaches a mismatch, inducing a conformational change in the protein such that the GIY-YIG endonuclease domain creates a double strand break in the DNA. This break is then repaired via homologous recombination (gene conversion). B) Hypothesized process (37) through which MSH1-induced gene conversion increases rate of heteroplasmic sorting in mitochondria. A heteroplasmic progenitor population contains two different alleles (green and blue). When MSH1 is present (left), it promotes gene conversion events (arrows), resulting in faster homogenization of gene copies within populations and increased variance among populations. In the absence of MSH1 (right) these gene conversion events are less common and variation in heteroplasmic frequencies accumulates more slowly.

Another way in which MSH1 could alter sorting dynamics is through changing the physical interactions and fusion events between mitochondria. Hypocotyl cells of *msh1* mutants show increased mitochondrial connectivity over wild type (85), suggesting an increased capacity for genome mixing and homogenization. Under existing models (14, 37), increased rates of fusion are predicted to accelerate mitochondrial sorting, the opposite of what we see in *msh1* mutants. However, if the lack of functional MSH1 results in lower rates of gene conversion, higher rates of mitochondrial fusion may not be sufficient to increase cell to cell variance. This observation, in combination with the finding that plastids sort faster than mitochondria even though they do not undergo fusion, suggests that the relationship between fusion and heteroplasmic sorting may be more complex than suggested by current models.

### Evidence for GC-biased gene conversion in plant organelles

Organellar genomes, including those of Arabidopsis (31, 86), are typically AT rich but also exhibit a wide range in nucleotide composition (58). The reasons for these biases are not fully understood, but most organisms across the tree of life experience an AT-biased mutation spectrum (87–89). However, in nuclear genomes, this mutation bias may be offset to varying extents by GC-biased gene conversion (90, 91). The possibility for biased gene conversion in plant organellar genomes is largely unexplored, and the few studies that have investigated this in plastids have come to differing conclusions about GC versus AT bias (92–94). In the SNVs studied here, we found evidence for GC-biased changes in allele frequency in both plastids and mitochondria, as well as in *msh1* and wild-type backgrounds. Notably, we observed this bias regardless of whether the mutant allele was AT (Mito 334038) or GC (Mito 91017, Plastid 26553, and Plastid 36873). Thus, it appears to be directly related to nucleotide composition and not a systematic preference for or against the reference allele. All four SNVs that were tracked in this study were in intergenic regions (Table 1), reducing the likelihood that the variants alter organellar or cellular function. In human mitochondria, variants in the non-coding D-loop did not show significant selection against pathogenic alleles; however, a broad population study showed a notable absence of variants in sites involved in gene transcription and mitochondrial DNA replication, in addition to other non-coding sites with no known function (10). Thus, a bias due to selection on functional effects of the variants studied here cannot be ruled out, especially given the incomplete characterization of regulatory elements and non-coding RNAs in plant organelle genomes (95–98).

If the observed GC bias is driven by gene conversion, it would suggest that the GC allele is favored during homologous recombination between heterogeneous DNA copies. In mitochondria, gene conversion could happen within heterogeneous organelles, but we expect it to be a particularly powerful actor during fusion in the SAM. In plastids, which do not undergo fusion, gene conversion is only expected to take place within organelles (not between organelles). This raises an important point about our sampling design. The SNVs we tracked are inferred to have arisen in the F2 generation, but we sampled mothers from later generations (F3 to F5 in plastids, Table S1). Therefore, we did not analyze the initial dynamics of plastid variants at their inception when they were present at low frequency among the genome copies within a single plastid. It is possible that some or even all of the plastids in our sampled mothers had already reached homogeneity for one allele or another, meaning that heteroplasmy dynamics were playing out among rather than within plastids. However, if the observed trend towards increasing frequency of GC alleles is the result of gene conversion bias, it would suggest that the mother plants in our study still contained plastids with copies of both alleles. This may be the case, but our data suggest extremely rapid heteroplasmic sorting, which might be expected if it were happening at the level of the small number of whole plastids within a cell. On the other hand, if the sets of genomes within each plastid had already reached homogeneity, it would suggest some unknown mechanism favoring GC alleles. These uncertainties emphasize the importance of determining how the rate of sorting among the genome copies within a plastid compares to the sorting process among the multiple plastids within a cell.

### Conclusions

The mechanisms of organellar genome maintenance and transmission are fundamentally different in plants and animals, but very little is known about how this affects the fate of variants arising from de novo mutations. We found that heteroplasmies in Arabidopsis organelles sort very rapidly which is likely due to a combination of low genome copy numbers in germline/progenitor cells and the recombinational nature of plant genomes. Our work supports a role for gene conversion as an important mechanism facilitating rapid sorting of heteroplasmic variants in Arabidopsis, which has been hypothesized to be a key mechanism for increasing variance in eukaryotic systems without early germline sequestration (37). Notably, the recombination surveillance and DNA repair gene *MSH1*, which is absent from most eukaryotic systems including animals (48), may play a key role in this process.

## Methods

### Plant material

Two homozygous *msh1* (At3g24320) mutant lines containing point mutations that result in either a nonsense mutation (CS3372: *chm1-1*) or an aberrant splice site (CS3246: *chm1-2*) were obtained from the Arabidopsis Biological Resource Center (50). Crossing design and techniques used for identification of de novo mutations are described previously (48). Briefly, homozygous mutants were used as males in crosses onto wild-type plants. The resultant F1 plants were allowed to self-pollinate, seed was planted and homozygous *msh1* lines (F2 families) were identified. The F2 lines were allowed to self-pollinate, and mitochondria and chloroplast were isolated from their progeny (pools of F3 plants). Organellar DNA was extracted and analyzed with Duplex Sequencing (99) to identify de novo mutations. High frequency SNVs identified in F3 organellar DNA pools from *msh1* lines were used in this study (Table 1).

In initial experiments, seeds were vernalized in water for 3 days at 4 °C, planted directly into 3-inch pots containing Pro-Mix BX media and grown on light shelves under short day conditions. Heteroplasmic individuals were moved to long (16 h light) day conditions when they began to bolt. Once siliques were ripe, all seeds were harvested in bulk. Seeds of heteroplasmic mothers from lines selected for further analysis were sterilized and plated on MS-agar (100), vernalized for 3 days at 4 °C and placed on light shelves to germinate. When seedlings had two true leaves, they were transferred to 1-inch pots filled with Pro-Mix BX media and placed in a growth chamber under short (10 h light) day conditions. Plants were transferred to long day conditions upon bolting.

### DNA extraction and heteroplasmy analysis

Tissue samples were disrupted using the TissueLyser (Qiagen), and total cellular DNA was extracted using the Qiagen Plant DNeasy kit. In cases where tissue was limiting (e.g., inflorescences), DNA was extracted by grinding tissue in 200 mM Tris-HCl pH 9.0, 250 mM NaCl, 25 mM EDTA, and 1% SDS, followed by precipitation in isopropanol. gDNA was quantified using Qubit (concentrations ranged from 0.5 - 30 ng/µL), and gDNA integrity of selected samples was checked by agarose gel electrophoresis.

Heteroplasmy analysis was performed with allele-specific ddPCR assays essentially as described in Wu et al. 2020 (48). Primers and probes for ddPCR were designed to ten different high-frequency SNV targets, seven in the plastid and three in the mitochondria (Table 1). Primers (Table S6) were designed to amplify fragments of 130 – 250 bp, with the SNV in the middle of the amplified sequence. Probes (Table S7) were designed to either the reference sequence or the variant sequence with the target SNV in the center. All primers and probes were synthesized by Integrative DNA Technologies.

ddPCR reactions were composed of: Bio-Rad ddPCR Super Mix for Probes (no dUTP), 250 nM final concentration of each (reference and variant) probe, 900 nM final concentration of each primer (F and R), 1 µL of the restriction enzyme BglII (which is used to fragment template DNA but is not predicted to cut within any of the amplified products), and 5 µL of an appropriate dilution of DNA in a 20 µL total reaction. Droplet generation was performed using a Bio-Rad QX200 Droplet Generator as per the manufacturer’s instructions, and PCR was performed in a Bio-Rad C1000 with a deep-well block under the following thermal cycling conditions: enzyme activation at 95 °C for 10 min, 40 cycles of 94 °C for 30 sec, annealing/extension temperature (see Table S6) for 1 min, and deactivation of the polymerase and restriction enzyme at 98 °C for 10 min – with a ramp speed of 2 °C per sec for all steps. Droplets were read on the Bio-Rad QX200 Droplet Reader and analyzed using QuantaSoft Analysis Software (Bio-Rad).

Dilutions of mutant (*msh1*) organellar DNA (48) were used to verify the presence of the SNV in the original mutant F3 organellar extractions alongside paired wild type samples to check for probe specificity. Wild type and mutant organellar samples and no template controls were used to determine appropriate annealing temperatures for each primer probe set to minimize off target binding in wild type and increase separation of positive and negative droplets in both channels. Positive and negative controls were always run alongside experimental samples to ensure assay fidelity and verify appropriate settings for channel thresholds. Background rates of the variant probe binding to wild type samples were typically very low < 0.05%. Experimental samples with variant calls falling at or under wild type values were set to zero for further analyses.

### Organellar genome copy number

Evagreen ddPCR was performed, essentially as described previously (101) to determine the number of mitochondrial and plastid genomes per nuclear genome copy. The reason for this was two-fold. First, it was important to determine whether the *msh1* mutant background altered the relative numbers of organellar genome copies. Secondly, numts (nuclear mitochondrial DNA, i.e., copies of mitochondrial DNA inserted into the nuclear genome) are known to be present in the *A. thaliana* nuclear genome (102–104) and we wanted to correct for this in our heteroplasmy analysis.

Two primer sets were used per genome to determine copy number (Table S1). Nuclear genome markers were single copy and located on chromosomes 1 and 2. Organellar markers were designed to single-copy mitochondrial (*rps12* and cox2) and plastid (*clpP1* and *psaA*) genes. For the nuclear genome, values from each primer set were averaged and used to calculate the number of organellar genome copies per nuclear genome for each organellar primer set. For plastids, values from each primer set were averaged to generate a final number of plastid copies per nuclear genome. For mitochondria, both primer sets amplify numts, so the values were adjusted based on the number of numt copies. The amplified regions of *cox2* and *rps12* are present at one and three numt copies, respectively, in the Arabidopsis nuclear genome (104). These values were subtracted from the calculated values of mitochondrial copies per genome before averaging to obtain a final value. Our mitochondrial SNVs of interest (mt91017 and mt334038) are both present in three copies in the Arabidopsis nuclear genome (104).

The number of organellar genomes per nuclear genome were determined for paired wild type and *msh1* mutants that were grown in parallel. Leaves were sampled when plants were 8 weeks old. For tissue specific analyses of mitochondrial genomes, samples consisted of whole inflorescences (n = 13) and 8-week-old leaf tissue: old leaves harvested from the base of the rosette (n = 12), and young leaves harvested from the top of the rosette (n = 16). A smaller subset of these was used to determine tissue specific amounts of plastid genomes. For each tissue, an average value was calculated for mitochondrial and plastid genomes per nuclear genome.

We used experimentally determined values of organellar genome copy number along with the deduced number of numt copies for each mitochondrial SNV (*Nu* = 3 in both instances) to correct our heteroplasmy values by computing a correction factor as described below:

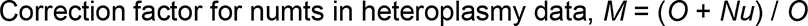

Where *O* = organellar genomes per nuclear genome copy, and *Nu* = nuclear genome copies of the SNV. The alternative allele frequency for a sample was multiplied by the correction factor *M*.

### Heteroplasmy sampling across generations

To understand the extent to which heteroplasmy is transmitted across generations, we identified heteroplasmic individuals in an initial screen and then analyzed the distribution of allele frequencies in their progeny using ddPCR. Initial screens to identify heteroplasmic mothers were conducted on F3 and/or F4 *msh1* mutant individuals. Leaves were sampled after 4-6 weeks of growth. Seeds from heteroplasmic individuals were sown and an initial leaf sample was taken from each offspring after 4 weeks of growth.

### Heteroplasmy sampling within plants

To understand how heteroplasmy is distributed within individuals, we selected three plants from each heteroplasmic mother (described above) for further tissue sampling. Three fully expanded rosette leaves were harvested from each plant at 5 weeks of growth. At 8 weeks of growth, leaves from the top (young) and base of the rosette (old) were harvested. Once plants began to bolt, entire single inflorescences were harvested. Selected tissue samples (8-week-old leaves and inflorescences) from this experiment were used to determine organellar genome copy number in experiments described above. This initial experiment did not lead to the identification of individuals heteroplasmic for plastid markers, so additional plastid SNV lines were grown and tissues were sampled in a subsequent experiment of the same design.

### Bottleneck calculations

We used two methods for estimating bottleneck size. First, we applied the common approach based on measures of heteroplasmic variance:

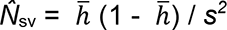

where *s^2^* is the variance of heteroplasmic frequency in progeny (or within a plant), (1/*n*) ∑ (*h_i_* – *h̄*)^2^, and *h̄* and *h_i_* are the sample mean and individual heteroplasmic frequencies of offspring (or tissues), respectively. In our intergenerational analysis, we used the offspring mean as an approximation of the maternal heteroplasmic frequency, even though we obtained experimental estimates of maternal heteroplasmy levels. This approach was chosen because we saw bias towards GC alleles (Figure 4), suggesting that maternal values would not be an effective estimate of average offspring values.

The second approach we took to estimate the bottleneck size was using a Kimura model (55). Here we maximized the joint likelihood of a set of heteroplasmy measurements under the Kimura model, which takes two parameters: mean heteroplasmy (*p*) and bottleneck parameter *b.* is related to the effective “bottleneck size” *N* by *N* = 1/(1-*b*). Previous work has proposed simply setting the population parameters to match the sample statistics *p* and *b* (55), but as this does not in general yield the maximum likelihood parameter estimates, it can give misleading results and does not support hypothesis testing. Instead, we used the kimura package in R (https://github.com/lbozhilova/kimura) to compute likelihoods and optimization using Nelder-Mead and Brent algorithms (105) to explicitly find the maximum likelihood parameters and the Fisher information matrix, from which we derive 95% confidence intervals. For homoplasmic cases, numerical issues challenged the Fisher approach, and bootstrap resampling with 200 resamples was instead used to estimate confidence intervals.

For hypothesis testing regarding the bottleneck size in two different groups of sample sets, we considered two statistical models. First, each set of samples is generated from the Kimura distribution with a set-specific *p* and a group-specific *b*. Second, each set of samples is generated with a set-specific *p* and a *b* common to both groups. For example, consider a comparison between mitochondrial and plastid intergenerational bottlenecks. In the first model, each family would have its own *p*, mitochondrial families would have one *b* value, and plastid families would have another *b* value. In the second model, each family would have its own *p* and a *b* common to all families. We then maximize the joint likelihood over all observations for the two models and conduct a likelihood ratio test with one degree of freedom, reflecting the additional *b* parameter in the first model. When reporting bottleneck size across samples in a given group (for example, across *msh1* mitochondrial families), we give the maximum likelihood estimate (*N̂*) and confidence intervals from this within-group inference. All code is freely available at https://github.com/StochasticBiology/plant-odna-sorting/.

### Impact of *MSH1* on heteroplasmy transmission

To determine whether MSH1 influences the spread of heteroplasmy across generations, we backcrossed *msh1/msh1* mutant females heteroplasmic for an SNV (either mt334038 or pt26553) to wild-type males. A minimum of 20 F1 plants were tested for heteroplasmy. Heteroplasmic plants were self-pollinated, and then F2 seedlings were planted and screened for heteroplasmy. Both F1 and F2 seedlings were genotyped at the *MSH1* locus as described in Wu et al. 2020 (48). F2 plants that were homozygous for the *msh1* mutant allele were removed from subsequent analyses.

## Acknowledgements

This work was supported by a grant from the National Institutes of Health (R01 GM118046 to DBS). This project has received funding from the European Research Council (ERC) under the European Union’s Horizon 2020 research and innovation programme (Grant agreement No. 805046 (EvoConBiO) to IGJ).

## Author contributions

AKB: Conceptualization, Investigation, Formal analysis, Validation, Visualization, Writing – Original Draft Preparation, Writing – Review and Editing

LK: Investigation, Formal analysis, Validation, Writing – Review and Editing

MFG: Investigation, Formal analysis, Writing – Review and Editing

MH: Investigation, Writing – Review and Editing

IGJ: Formal analysis, Funding Acquisition, Validation, Visualization, Writing – Review and Editing

DBS: Conceptualization, Formal analysis, Funding Acquisition, Visualization, Writing – Original Draft Preparation, Writing – Review and Editing

## Supporting Information

**Figure S1.**
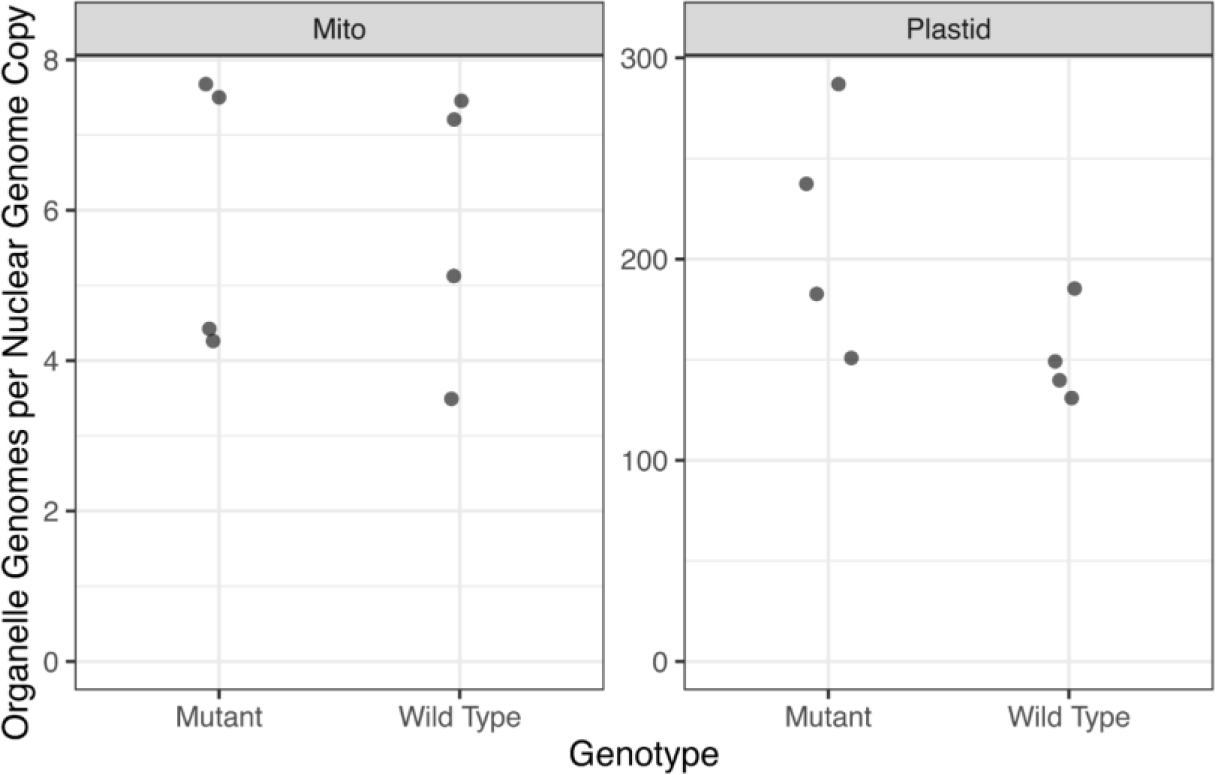
MSH1 does not significantly alter organellar genome copy number. The number of mitochondrial and plastid genome copies per nuclear genome copy was determined using ddPCR, corrected for numts and compared between *msh1* mutant lines (n = 4) and wild type plants (n = 4). Organellar genome copies between mutant and wild type lines were not significantly different (mitochondria, p = 0.910; plastid, p = 0.098).

**Figure S2.**
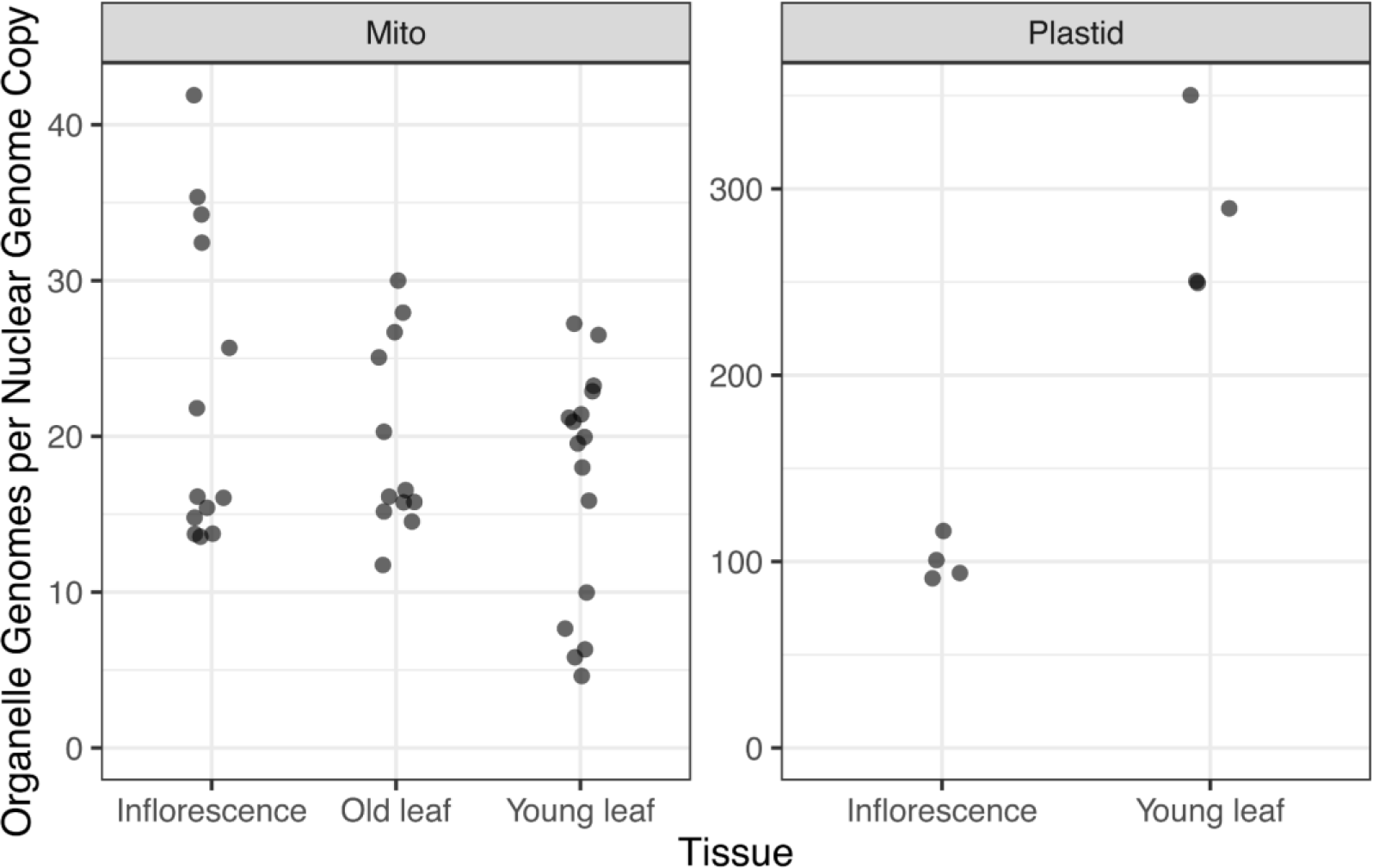
Organellar genome copy number varies within and between tissues. The number of mitochondrial and plastid genome copies per nuclear genome copy was determined using ddPCR and corrected for numts. For mitochondrial primers, tissue samples consisted of whole inflorescences (n = 13), old leaves harvested from the base of the rosette (n = 12), and young leaves harvested from the top of the rosette (n = 16). A subset of samples was chosen for analysis using plastid primers (inflorescences, n = 6; young leaf, n = 6). Plastid copies were significantly lower in inflorescences versus leaves (p = 0.00028); mitochondrial copies were not significantly different between tissue types. Mitochondrial values were adjusted for numt copies.

Table S1. Distribution of SNVs over multiple generations of *msh1* mutant lines.

Table S2. Values of heteroplasmy for mother and progeny.

Table S3. Within plant heteroplasmy.

Table S4. Distribution of SNVs over two generations in wild type background.

Table S5. GC bias.

Table S6. Primers for ddPCR heteroplasmy assays and genome copy analysis.

Table S7. Allele-specific probes for ddPCR heteroplasmy assays.

